# A Procoagulant Peptide Analog of the SARS-CoV-2 Nucleocapsid C-terminal Domain

**DOI:** 10.64898/2026.07.04.736500

**Authors:** Brian Andrich L. Pollo, Fresthel Monica M. Climacosa, Salvador Eugenio C. Caoili

**Affiliations:** Biomedical Innovations Research for Translational Health Science (BIRTHS) Laboratory, Department of Biochemistry and Molecular Biology, College of Medicine, University of the Philippines Manila, Manila, Philippines; Department of Medical Microbiology, College of Public Health, University of the Philippines Manila, Manila, Philippines

**Keywords:** Hemostasis, Peptides, Blood coagulation, Blood plasma, Blood coagulation factors

## Abstract

**Background:** Uncontrolled bleeding complicates trauma, surgery and many medical conditions. While currently available procoagulant therapies (*e*.*g*., plasma-derived factors, recombinant proteins, antifibrinolytics) have crucial limitations.

**Methods:** N389 (CQQTVTLLPAADLDDFSC) was synthesized by Fmoc solid-phase chemistry, characterized by HPLC and LC–MS, then tested in normal human pooled plasma in microplate mechanical clot-formation assays using incubated and immediate addition formats. Kinetic parameters (plasma recalcification, PRT; maximum absorbance, A_*max*_) were obtained from absorbance curves fit to four-parameter logistic models. Mixing studies with modified (*i*.*e*., aged, adsorbed) plasma probed factor dependence.

**Results:** In plasma coagulation assays activated with 25 mM CaCl□, baseline clotting showed a PRT of 23.74 ± 0.27 min and A_*max*_ of 0.1813 ± 0.0043 (*n* = 3), whereas N389 significantly reduced PRT to 8.442 ± 6.0395 min without incubation (*p* = 0.0012), further decreased PRT after incubation (*p* < 0.0001), increased A_*max*_ to 0.2523, and retained comparable activity across normal, adsorbed, and aged plasma, in contrast to S1255 which showed a faster but incubation-labile effect with PRT 2.353 ± 1.3685 min (*p* = 0.0007) and marked attenuation in factor-depleted and aged plasma. Mixing studies showed N389 activity persisted across normal, aged and adsorbed plasma, consistent with a mechanism that does not require intact plasma coagulation factor profiles (specifically factors II, V, VIII, VII, IX, X).

**Discussion:** Collectively with prior evidence on anionic surfaces, Ca^2^□-binding Gla domains, and peptide-modulated fibrin polymerization, these results support a model in which N389 functions as a stable, charge-based scaffold that coordinates divalent cations and/or directly nucleates fibrin(ogen), while highlighting limitations of bulk clotting assays and the need for targeted thrombin generation, binding, aggregation, and contact-activation studies.

**Conclusions:** The aspartate-rich peptide N389 is a sustained, factor-independent procoagulant at least *in vitro*. N389 thus merits further mechanistic and translational evaluation as a synthetic hemostatic agent.

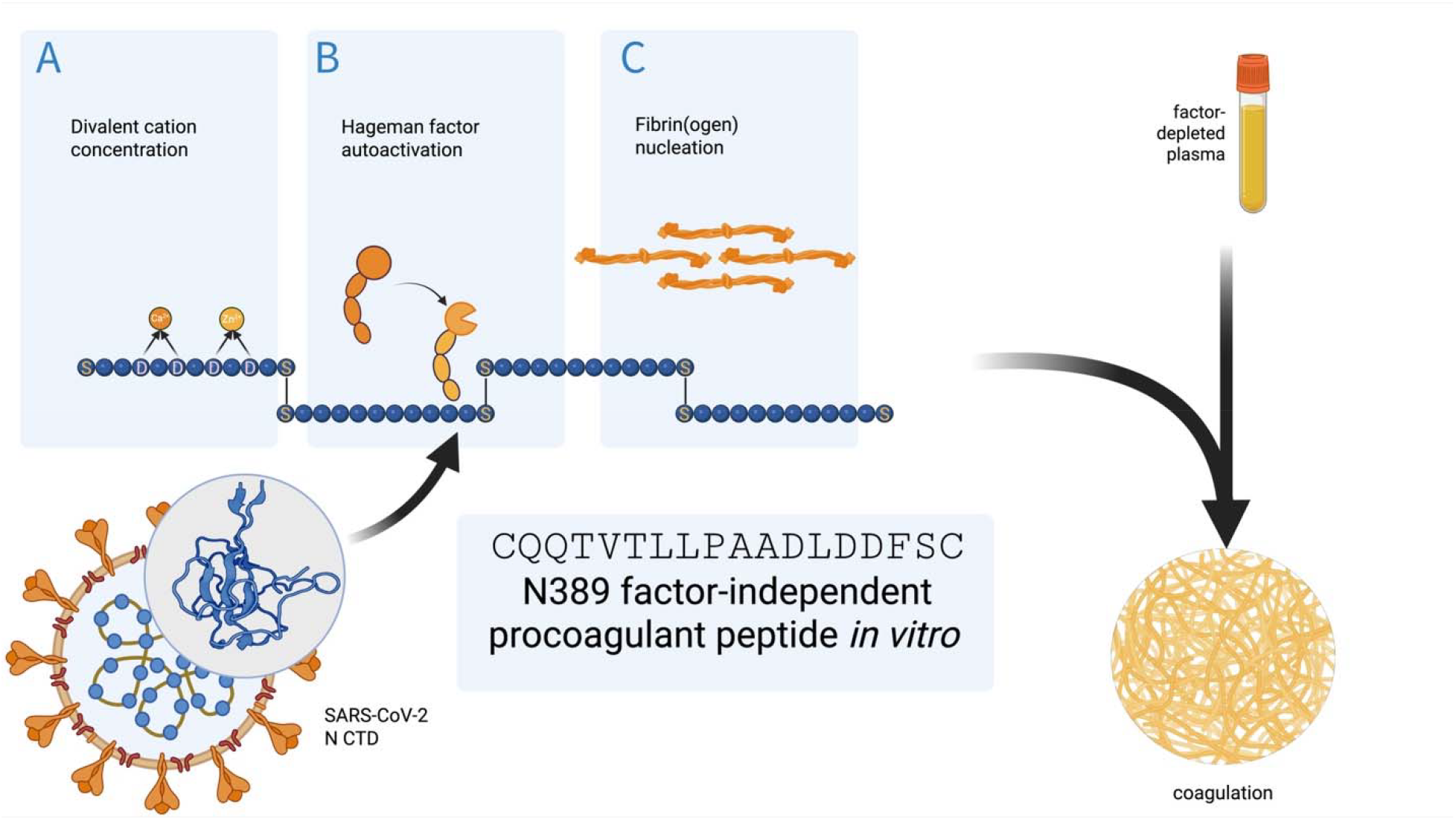

## INTRODUCTION

Coagulation is a vital physiological process that prevents hemorrhage following vascular injury, relying on the tightly regulated interplay of cellular and plasma factors within the coagulation cascade. Disruption of this balance whether through trauma, surgery, or congenital deficiencies, can lead to life-threatening bleeding diatheses that demand timely therapeutic intervention [1].

Current procoagulant strategies predominantly include plasma-derived clotting factor concentrates, recombinant proteins such as activated Factor VII, and antifibrinolytics like tranexamic acid. While effective in certain contexts, these agents present key limitations: whole-protein preparations carry structural complexity, batch variability, and potential immunogenicity; while recombinant proteins are costly, require cold-chain storage, and can elicit neutralizing antibodies, particularly in hemophilia patients with inhibitors [2–5]. Moreover, these therapies often exert broad-spectrum activity without the ability to selectively target specific points in the cascade, increasing the risk of off-target effects.

Synthetic peptides have emerged as a promising alternative, offering modular design, chemical definability, and batch-to-batch reproducibility [4,6]. Their small size and tunable sequences allow for precision in targeting, while chemical synthesis eliminates the risk of blood-borne pathogen transmission. Post-synthetic modifications can further improve stability, bioavailability, and specificity, attributes difficult to achieve with plasma-derived or recombinant protein therapeutics.

However, there is a scarcity of synthetic peptide candidates with potent, specific, and predictable procoagulant activity validated in clinically relevant models. This shortfall has hindered their translation into routine medical use, despite their theoretical advantages.

The present study addresses this gap by introducing an aspartate-rich peptide, perfectly homologous to a sequence within the SARS-CoV-2 nucleocapsid C-terminal domain, designed to enhance blood coagulation. The peptide is hypothesized to provide advantages in safety, manufacturability, and functional control over existing plasma-derived and recombinant protein-based procoagulants, potentially advancing the development of next-generation hemostatic agents [7,8].

## METHODS

### Peptide Synthesis and Characterization

The aspartate-rich peptide N389 was synthesized by a commercial provider (GenScript, Singapore) using Fmoc solid-phase chemistry. Peptide identity and purity (>95%) were confirmed by LC–MS (ESI positive-ion mode, scan range 300–2000 m/z). Analytical HPLC employed an Inertsil ODS-3 column (4.6 × 250 mm; GL Sciences, Tokyo, Japan) with solvent A (0.065% TFA in water) and solvent B (0.05% TFA in acetonitrile), flow rate 1.0 mL/min, gradient from 5% to 65% B over 25 min, UV detection continuous at 220 nm. Lyophilized peptide was reconstituted in sterile deionized water, aliquoted, and stored at −20 °C. For oxidative polymerization, peptides were incubated in 50% (v/v) DMSO at 25 °C for 24 h.

### Incubated Coagulation Assay

Pooled normal human plasma (from 25 healthy donors, fresh frozen plasma provided by the blood bank of the Philippine General Hospital) was thawed at room temperature and equilibrated to room temperature. In 96-well plates, 30 µL plasma was incubated with peptide (20 µg/mL final concentration) or control (buffer only) at 37 °C for 1 h. Then, 60 µL HEPES-buffered saline (HBS; 20 mM 4-(2-hydroxyethyl)-1-piperazineethanesulfonic acid, 150 mM NaCl, pH 7.4; Thermo Fisher) was added. Clotting was initiated with 30 µL of 100 mM CaCl□(25 mM final) delivered using a multichannel pipette to standardize the start time. Plates were transferred into a prewarmed (37 °C) microplate reader (BioTek), and absorbance at 450 nm was recorded every 5 min for 60 min, starting at 2 min. Clot presence was confirmed post-run by adding 5 µL dye solution (0.12% trypan blue, 10% glycerol) per well, observing whether a surface clot formed.

Primary kinetic parameters were prothrombin reaction time (PRT; defined as time to half-maximal turbidity) and maximum absorbance (A_*max*_), derived by fitting the absorbance curve to a four-parameter logistic (4PL) regression model in GraphPad Prism 9 (GraphPad Software, San Diego, CA, USA). All assays were performed in triplicate (*n* = 3 biological replicates).

### Immediate Coagulation Assay

Peptide-treated plasma (20 µg/mL) was combined with HBS and immediately recalcified to 25 mM CaCl□, omitting the incubation period. Kinetic data were recorded as described above to evaluate time-dependent activity differences.

### Mixing Studies

To delineate factor dependence, mixing experiments were performed using normal, aged, and adsorbed plasma. Aged plasma (deficient in factors V and VIII) was prepared by storing pooled plasma at 37 °C for 24 h. Adsorbed plasma (deficient in Gla-factors II, VII, IX, X) was obtained by incubating plasma with 100 mg/mL BaSO□ for 15 min at 37 °C, then centrifuging at 500 × *g* for 3 min and collecting the supernatant.

For these assays, 15 µL plasma was incubated with peptide (20 µg/mL) in HBS (1 h, 37 °C), followed by addition of 30 µL modified plasma (normal, aged, or adsorbed) and dilution with 30 µL HBS. The mixture was recalcified and clotting kinetics recorded as in the incubated assay.

### Statistical Analysis

Data are reported as mean ± standard deviation (SD). Comparisons of kinetic parameters between groups (*e*.*g*., N389 versus control) were made using two-tailed unpaired *t*-tests when comparing two groups. A *p*-value < 0.05 was considered statistically significant. Statistical analyses were conducted using GraphPad Prism 9.

## RESULTS

Synthesis of the aspartate-rich peptide N389, homologous to a segment of the SARS-CoV-2 nucleocapsid carboxy-terminal domain, yielded a single major product (Supplementary Figure S1). LC–MS analysis identified a dominant peak at 18.640 min, representing 32.039% of the integrated peak area (Supplementary Figure S2). The mass spectrum of this peak displayed a principal ion at *m/z* 970.6 [M+2H]2+ (Supplementary Figure S3). Deconvolution gave an estimated molecular mass of 1939.18 Da, closely matching the predicted mass of 1939.2 Da, confirming successful synthesis and purity of the peptide.

In baseline coagulation assays activated by 25 mM CaCl□, clotting followed a sigmoidal kinetic profile, with a peak reaction time (PRT) of 23.74 ± 0.27 min and a maximal absorbance (*A*_max_) of 0.1813 ± 0.0043 (n = 3). The addition of N389 markedly accelerated clot formation, reducing PRT to well below baseline (*p* < 0.0001) and increasing *A*_max_ to 0.2523 (Figure 1). This indicated that N389 not only hastened clot initiation but also enhanced the extent of clot formation under incubated conditions.

**Figure 1.**
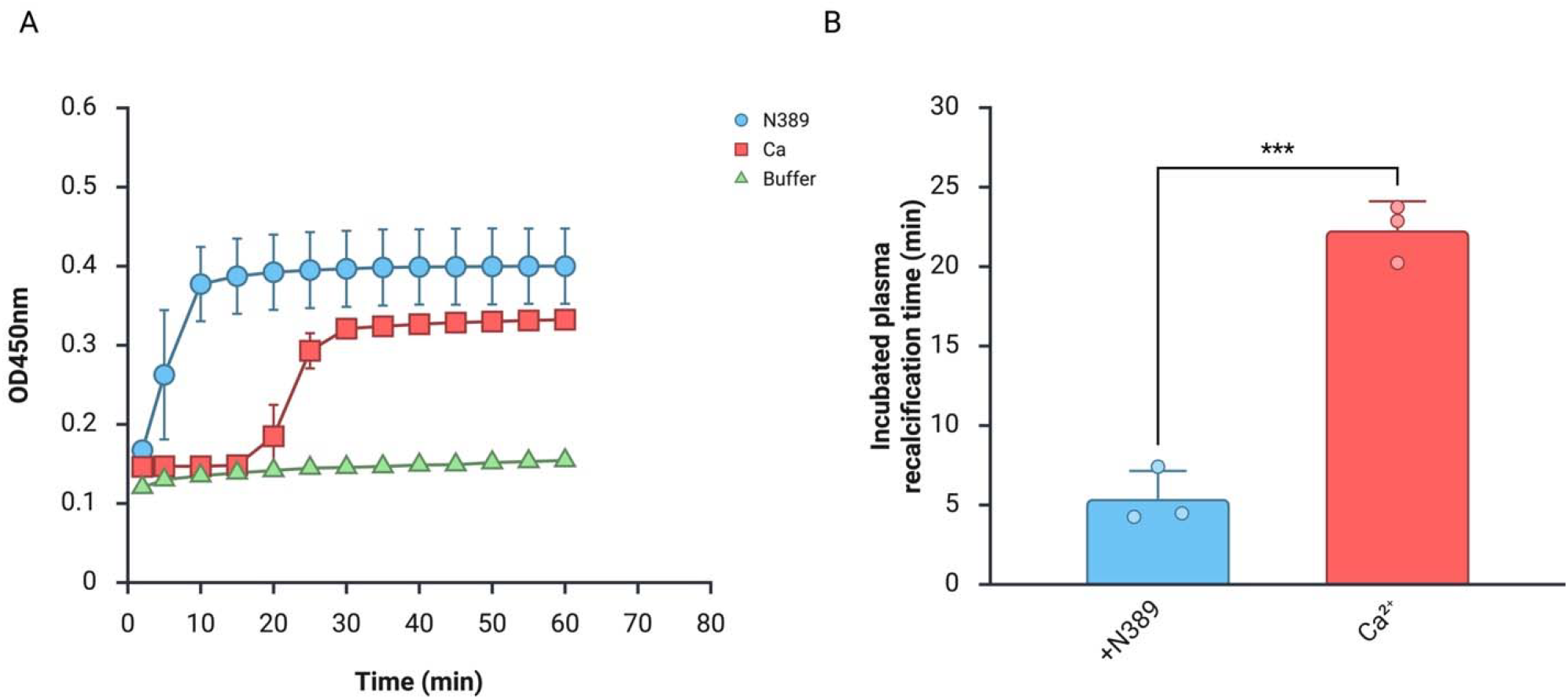
Incubation with N389 enhances coagulation activity of recalcified pooled normal human plasma. (A) Time series of clot formation along with (B) the plasma recalcification time from the data of (A) is shown. Data represent mean ± SD (*n* = 3); Paired *t-*test: *** p ≤ 0.001.

When tested without incubation, N389 continued to exert a strong procoagulant effect, shortening PRT to 8.442 ± 6.0395 min (*p* = 0.0012) (Figure 2). In comparison, the peptide S1255 produced an even more rapid onset of clotting, with a PRT of 2.353 ± 1.3685 min (*p* = 0.0007). The sustained activity of N389 after incubation, contrasted with the more transient effect of S1255, suggests that the two peptides act via distinct mechanisms.

**Figure 2.**
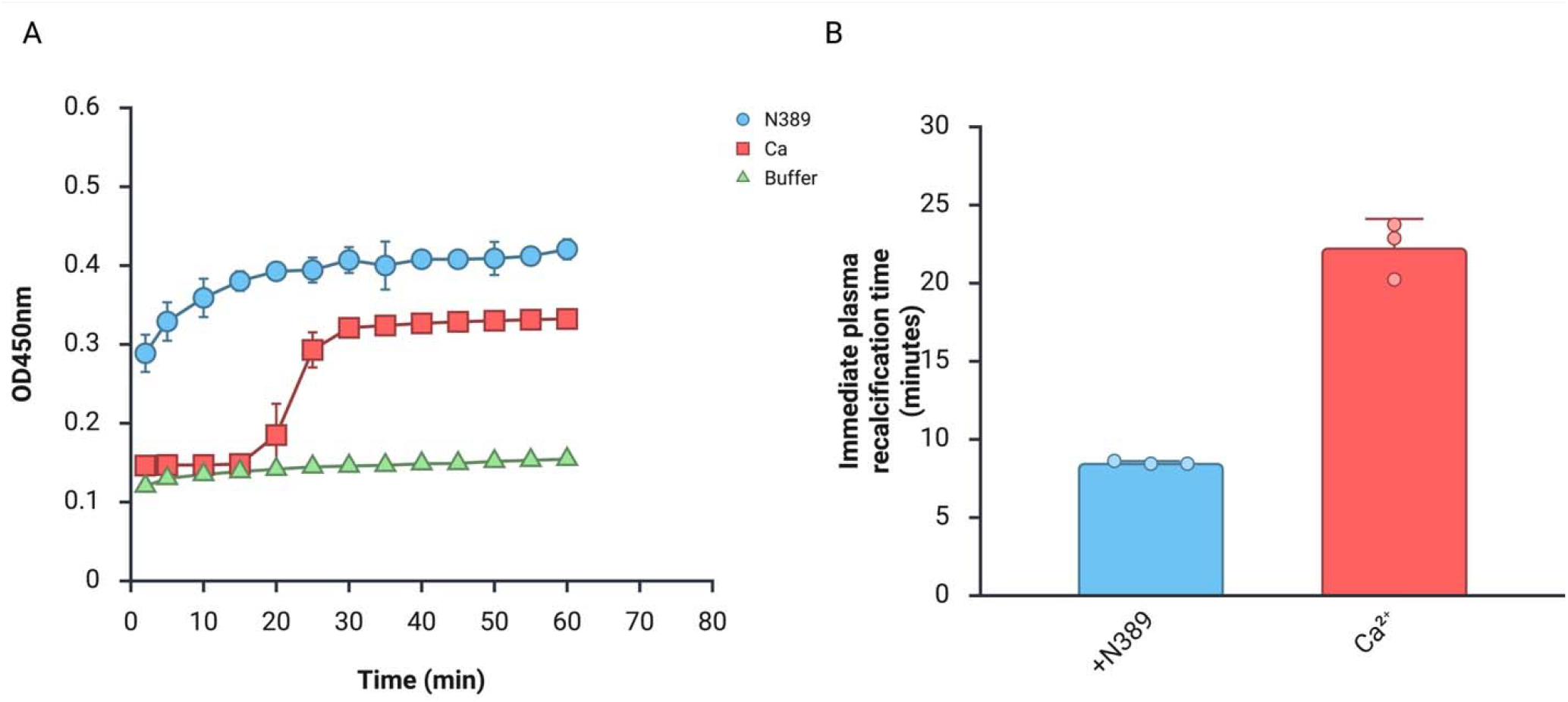
Immediate addition of N389 enhances coagulation activity of recalcified pooled normal human plasma. (A) Time series of clot formation along with (B) the plasma recalcification time from the data of (A) is shown. Data represent mean ± SD (*n* = 3); Paired *t-* test: * p ≤ 0.05.

Across normal, adsorbed, and aged plasma, N389 maintained a consistent procoagulant effect (Figure 3), implying that its activity is largely independent of the presence or integrity of plasma coagulation factors. In contrast, S1255 exhibited its highest activity in normal plasma, was attenuated in adsorbed plasma, and further reduced in aged plasma, indicating a greater dependence on intact plasma components for its procoagulant activity.

**Figure 3.**
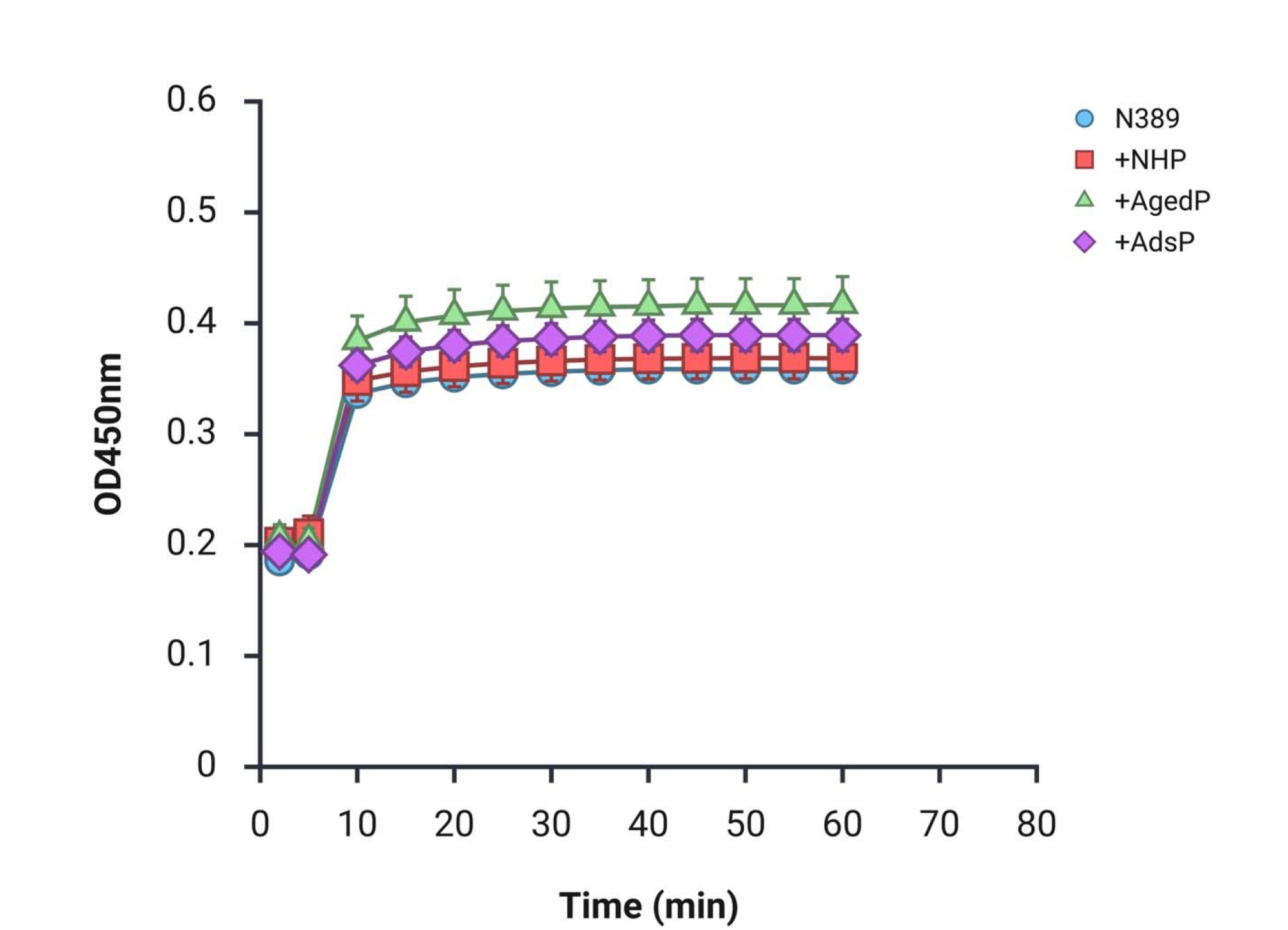
Coagulation curve of mixing studies of pooled normal human plasma incubated with N389, respectively, each mixed 1:1 with normal human plasma, aged plasma, or adsorbed plasma. Data represent mean ± SD (*n* = 3)

**Figure 4.**
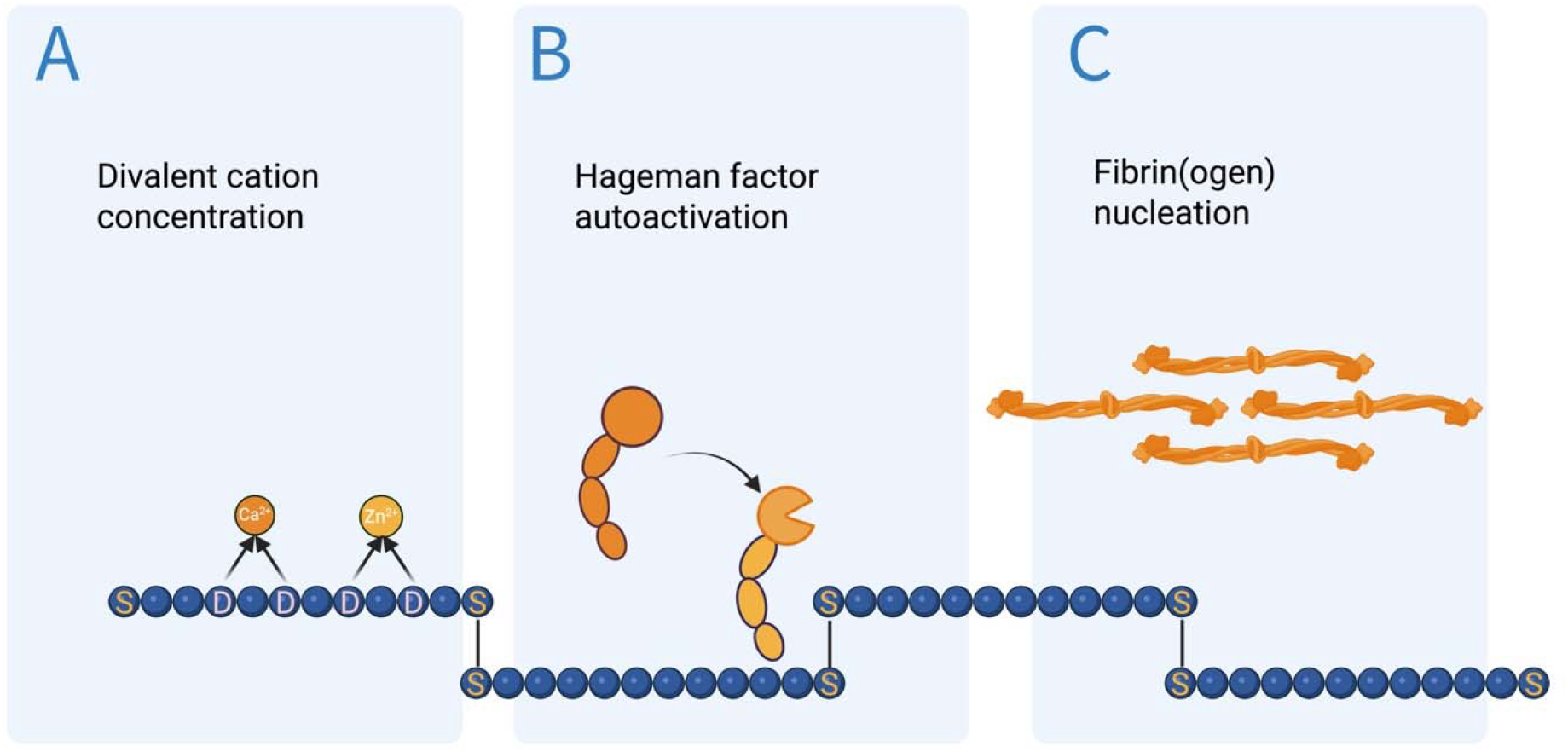

## DISCUSSION

The aspartate-rich peptide N389 produced an immediate procoagulant effect when added directly to plasma and retained a sustained procoagulant activity after incubation, whereas the comparator peptide S1255 produced a rapid but labile effect that diminished with incubation. Mixing studies further indicated that N389’s activity was preserved across normal, adsorbed, and aged plasma, consistent with an effect that does not strictly depend on intact levels of most soluble coagulation factors. From these observations we hypothesize that N389 promotes clot formation through a charge-based, surface-like mechanism, likely involving coordination of divalent cations (*e*.*g*., Ca^2^□and/or Zn^2^□) and/or direct interactions with fibrin(ogen), rather than through classical enzymatic amplification of specific clotting factors.

Several lines of prior work provide precedent for charge- or surface-mediated acceleration of coagulation and help situate our hypothesis. Biological polyanions such as inorganic polyphosphate accelerate contact activation and downstream thrombin generation by serving as an anionic surface that promotes assembly and activation of contact system components. Polyphosphate-mediated effects on FXII and thrombin generation have been demonstrated in both biochemical and in vivo models [9]. In the vitamin K–dependent coagulation factors, calcium binding to clusters of γ-carboxylated glutamate residues (Gla domains) is a well-established molecular mechanism that mediates membrane association and spatial organization of prothrombinase/tenase complexes. This provides a known example of how negatively charged/ion-coordinating motifs can position protease–substrate interactions for efficient thrombin generation [10]. Finally, the physical chemistry of fibrin polymerization, the knob-hole interactions and protofibril assembly that determine clot structure, can be modulated by exogenous peptides and scaffolds that alter local concentration, orientation, or kinetics of fibrin monomers [11]. Together, these literatures show that non-enzymatic, surface- or ion-mediated mechanisms can meaningfully accelerate clot formation and therefore are congruent with our working hypothesis for N389.

Mechanistically, several non-mutually exclusive models could explain the pattern of results observed. First, the dense array of aspartate side chains in N389 could act as a multivalent carboxylate surface that binds Ca^2^□(and possibly Zn^2^□) and thereby functions as a synthetic, peptide-based analog of the negatively charged membrane surface normally provided by phosphatidylserine; by localizing divalent cations, such a scaffold could enhance local effective concentration of ions required for factor–membrane or factor–factor interactions and accelerate formation of active complexes. This idea is supported conceptually by the centrality of Ca^2^□– Gla interactions in native coagulation factor membrane binding [10]. Second, the peptide could interact directly with fibrinogen or nascent fibrin, serving as a nucleation site that lowers the energy barrier for protofibril formation and accelerates polymerization; the literature on peptide-modulated fibrin assembly and on synthetic hemostatic peptides/materials shows that exogenous scaffolds can both speed clot formation and alter clot architecture [11,12]. Third, N389 could promote contact activation (FXII pathway) by presenting an anionic surface that supports FXII autoactivation or kallikrein amplification analogous to the action of long-chain polyphosphates; such a route could produce thrombin generation without relying on pre-existing high concentrations of other factors, consistent with the mixing-study results [9]. Any of these mechanisms, ion coordination, fibrin nucleation, or contact pathway facilitation, could account for an immediate effect and a form of sustained activity if the peptide forms stable complexes or aggregates that persist during incubation.

### Study limitations

There are important limitations to our study and to the mechanistic inferences that can be drawn from the present data. First, our coagulation assays relied on bulk clotting readouts (PRT, Amax); these do not directly measure thrombin generation, factor activation states, peptide–protein binding kinetics, or ion-binding stoichiometry. Second, the precise chemical status and aggregation state of N389 under assay conditions were not characterized beyond LC-MS of the synthesized product; peptide oligomerization or formation of calcium-rich nanoparticulates could be central to activity but remain unmeasured. Third, although mixing studies support factor-independence, they do not exclude involvement of specific plasma components (*e*.*g*., contact factors, high-molecular-weight kininogen, or serum proteins) that survive adsorption or aging differently. Fourth, the experiments reported here are entirely in vitro; in vivo hemostatic efficacy, potential prothrombotic risk, pharmacokinetics, and immunogenicity are unknown. Finally, the comparator peptide S1255’s behavior suggests different kinetics or stability, but without biophysical data (*e*.*g*., aggregation propensity, proteolytic susceptibility) definitive reasons for its loss-of-function after incubation cannot be ascertained.

Our results indicate that an aspartate-dense peptide can exert robust procoagulant effects in standard plasma assays and thus may represent a class of peptide-based hemostatic agents with potential utility in topical or biomaterial applications. The observation that plant-derived or synthetic peptides can influence coagulation is supported by prior reports of plant-derived peptide fragments and engineered self-assembling peptides that affect clotting or improve hemostasis in ex vivo and animal models, suggesting translational promise if safety and delivery challenges are addressed [13]. Nonetheless, differences in plasma composition, species, and wound microenvironment may modulate efficacy: peptides that act as insoluble nucleation scaffolds in a controlled in vitro setting could behave differently in the presence of cells, flow, proteases, or in the complex milieu of a bleeding wound.

We therefore recommend the following experiments to validate mechanism and advance translational evaluation. First, perform time-resolved thrombin generation assays to quantify lag phase, peak thrombin, and endogenous thrombin potential in the presence and absence of N389, and test sensitivity of these parameters to calcium chelation (e.g., EDTA) to probe Ca^2^□dependence. Second, carry out fibrin polymerization assays (turbidity and imaging) and scanning or transmission electron microscopy to visualize whether N389 alters fibrin fiber nucleation, branching, or network architecture. Third, use surface plasmon resonance or isothermal titration calorimetry to measure direct binding between N389 and fibrinogen, prothrombin, factor X/Xa, or divalent cations, and apply SPR to compare binding kinetics with S1255. Fourth, evaluate contact activation specifically (FXII activation assays, prekallikrein cleavage) to determine whether anionic surface-dependent pathways contribute. Fifth, characterize peptide aggregation and particulate formation under assay conditions (dynamic light scattering, nanoparticle tracking), since particulate formation may underpin a stable procoagulant surface. Finally, if these mechanistic studies support a safe, factor-independent procoagulant profile, progress to controlled *ex vivo* whole-blood and small-animal bleeding models alongside toxicity and immunogenicity assessments.

## CONCLUSION

N389 is an aspartate-rich peptide that produces immediate and sustained acceleration of clot formation *in vitro* and retains activity across varied plasma preparations. The pattern of results is most consistent with a charge- and ion-mediated, surface-like mechanism, potentially involving calcium coordination, fibrin nucleation, or contact pathway facilitation, rather than a purely factor-dependent enzymatic activation. Targeted biochemical and biophysical studies are now required to distinguish among these models and to determine whether N389 or related peptides can be developed into safe and effective hemostatic agents.

## ETHICS

Ethics approval was obtained from the University of the Philippines Manila Research Ethics Board (Code 2022-0492-01).

## FUNDING

This work was financially supported in part by dissertation grants from the Department of Science and Technology Philippine Council for Health Research and Development (DOST-PCHRD).

## CONFLICT OF INTEREST

The authors declare no potential conflict of interest with respect to authorship, and publication of this article.

## DATA AVAILABILITY STATEMENT

Supplementary figures and raw data are hosted in Harvard Dataverse (https://doi.org/10.7910/DVN/2M6QSW).

---

**Supplementary Figure 1.**
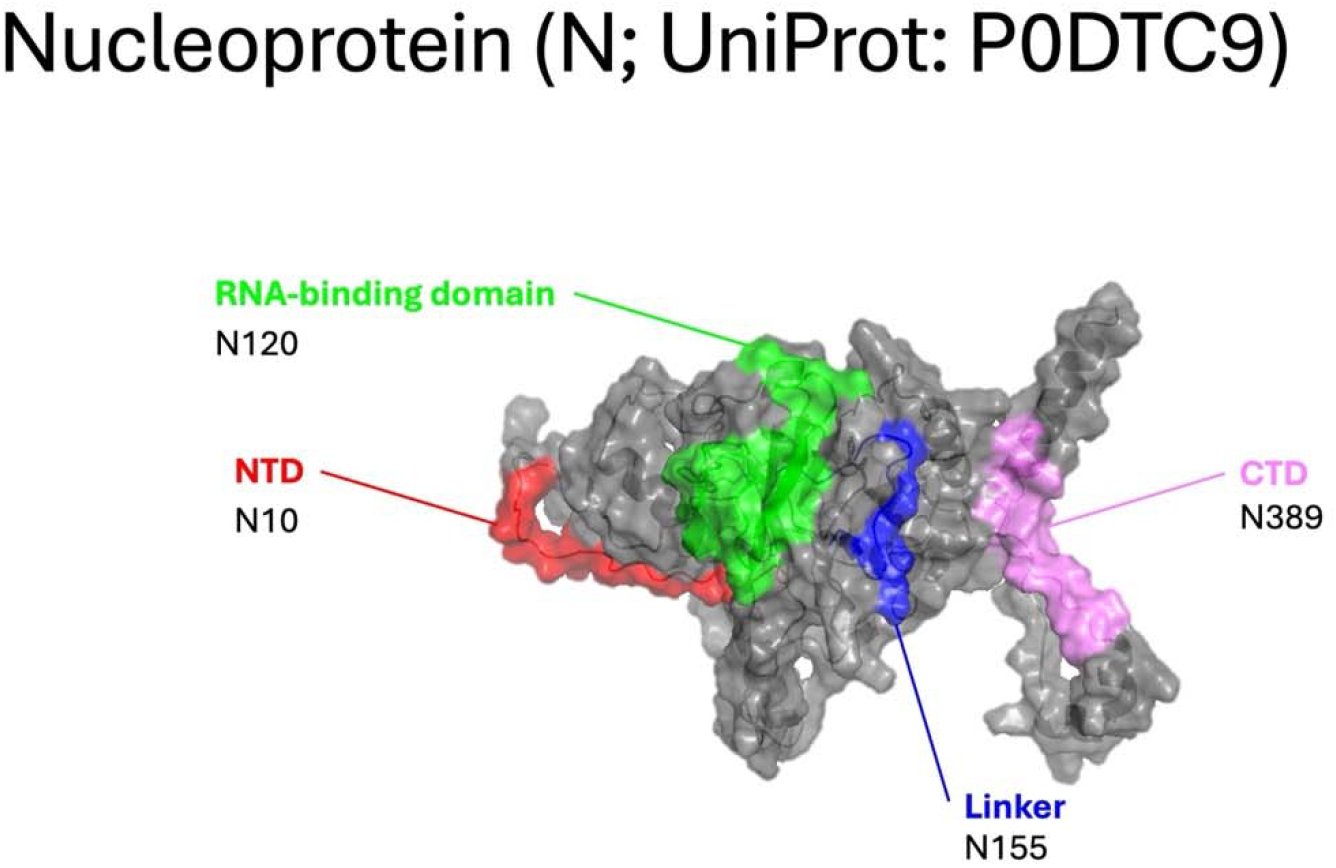
Space-filling molecular model of SARS-CoV-2 nucleocapsid protein, highlighting clusters of accessible, disordered linear segments; N389 was synthesized as an analog of the carboxy-terminal cluster.

**Supplementary Figure 2.**
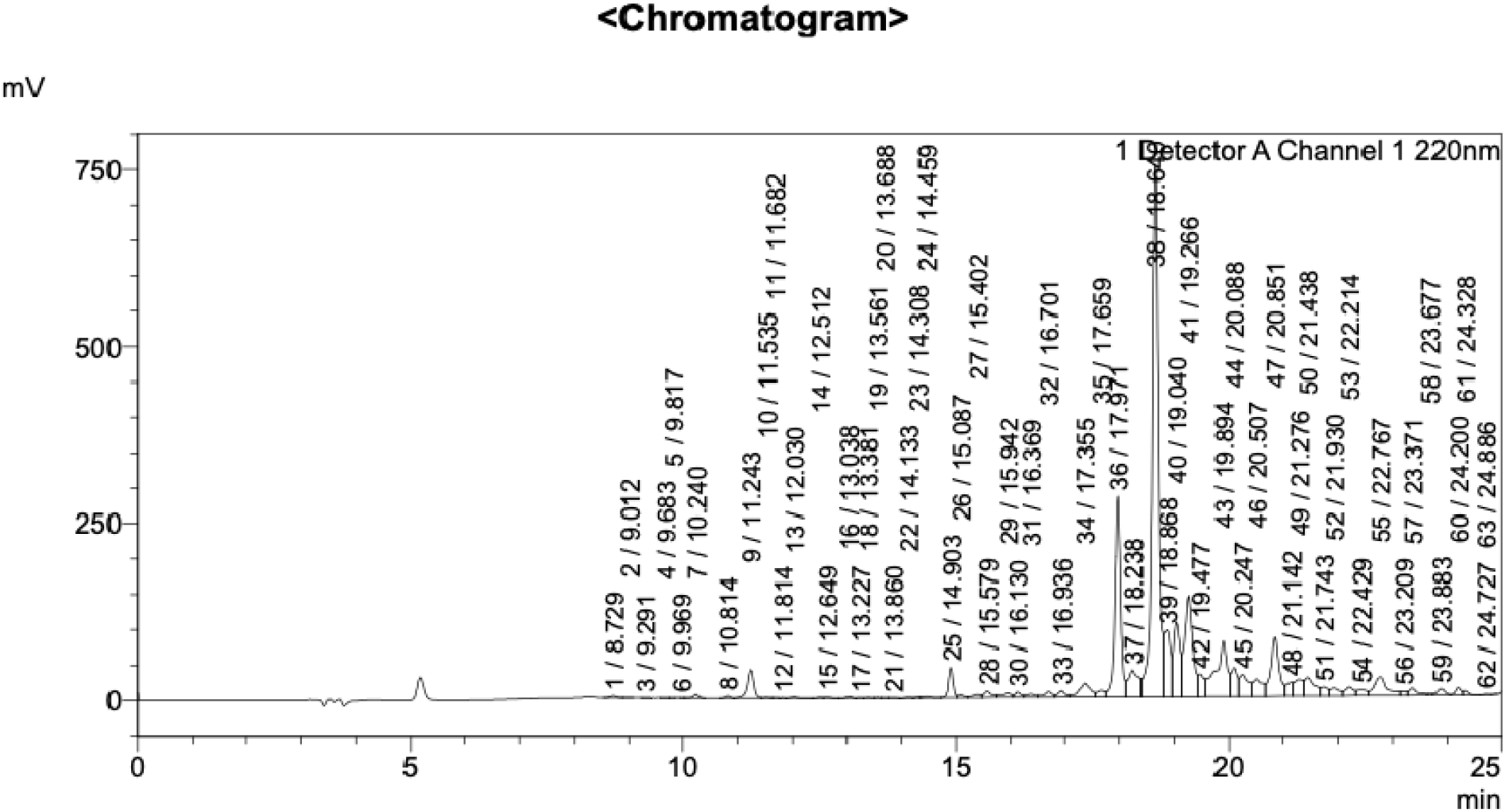
Liquid chromatography profile wherein the dominant peak, exhibiting an elution time of 18.640 minutes and comprising 32.039% of the integrated peak area, corresponds to N389, the synthetic peptide of sequence CQQTVTLLPAADLDDFSC.

**Supplementary Figure 3.**
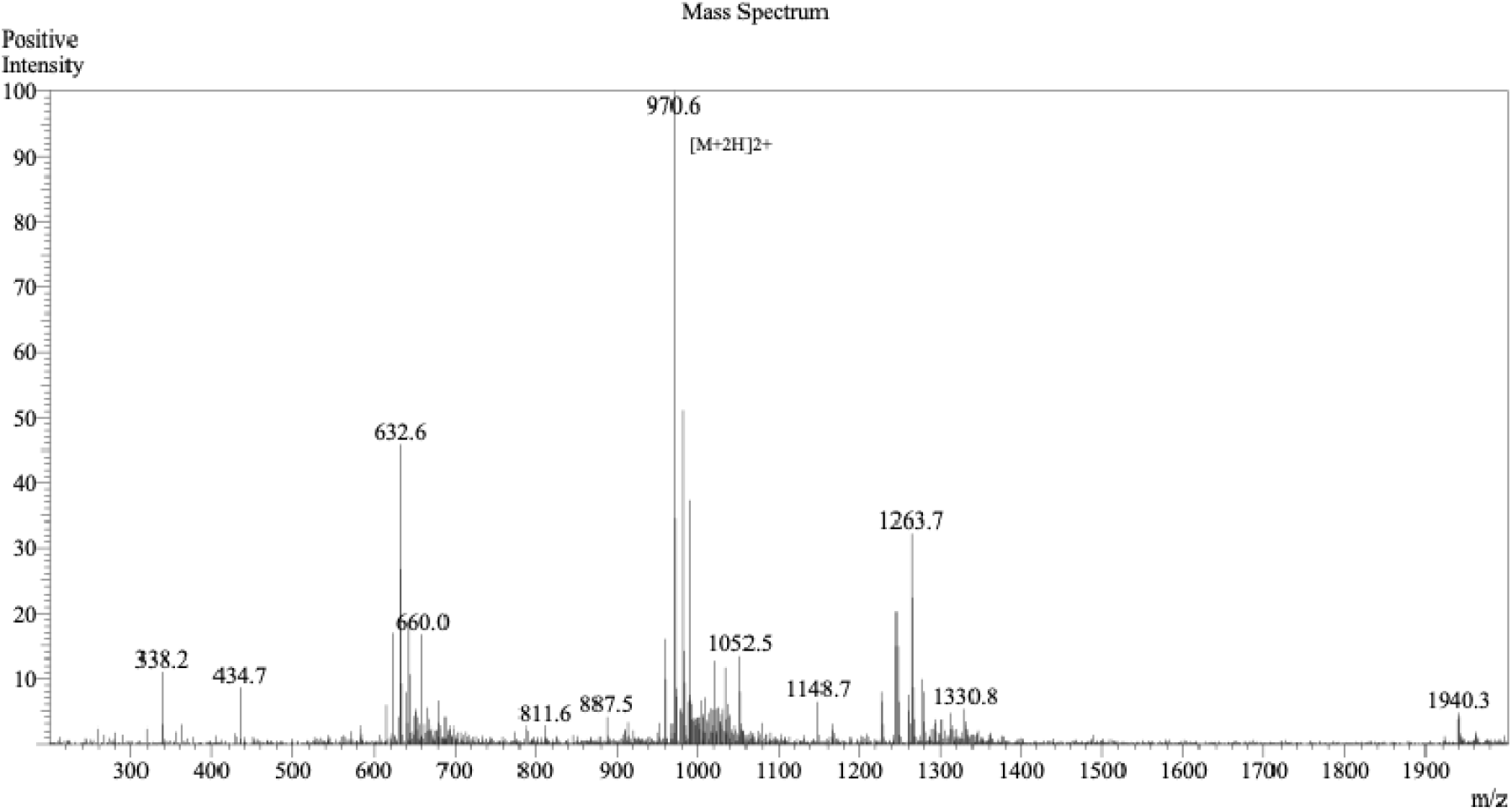
Mass-to-charge ratio (m/z) spectrum wherein the dominant ions correspond to N389, the synthetic peptide of sequence CQQTVTLLPAADLDDFSC.

## Notes

https://doi.org/10.7910/DVN/2M6QSW

### Competing Interest Statement

The authors have declared no competing interest.

